# Hypertension and diabetes mellitus association to oxidative stress in an elderly population

**DOI:** 10.1101/269993

**Authors:** Aleida Rodríguez-Castañenda, Katia Leticia Martínez-Gonzáles, Rosalinda Sánchez-Arenas, Sergio Sánchez-García, Israel Grijalva, Lourdes Basurto, Juan Cuadros-Moreno, Eliseo Ramírez-Garcia, Paola Garcia-delaTorre

## Abstract

**Background:** Mexico City has the highest aging rate in the country, as well as a high prevalence of diabetes mellitus (DM) and arterial hypertension (HT). All three on their own, are known to increase oxidative stress (OE).

**Methods:** Final groups included 18 patients without DM or HT (control group), 12 with DM, 23 with HT, and 18 with DM and HT. The EO was measured by the quantification of reactive oxygen species (ROS), and by determination of lipid peroxidation.

**Results:** HAS patients showed increased ROS levels as did men with HAS compared with the respective DM and HT groups. Also, women of the control group showed higher levels of ROS compared with men. HT in an aged population turned out to be the most influential factor for oxidative stress increase while DM had no effect whatsoever.

## Introduction

Aging is a gradual and adaptive process characterized by a diminished homeostatic response due to morphological, physiological, biochemical, and psychological modifications due to an accumulated deterioration, one of the reflections of this process is a vulnerability to chronic-degenerative diseases^1^.

World population is aging at an accelerated pace, this phenomenon refers to a growing proportion of people older than 60 years of age compared to any other group of age^2^. By 2015, Mexico had 1 elder (older than 60 years of age) per every 10 young adults (15 years of age), a number that will double by 2050^3^. Currently, Mexico City is the entity with the highest aging index with 61.7 elders per every 100 individuals^3^.

Diabetes Mellitus (DM) and arterial hypertension (HT) are the chronic-degenerative diseases that cause the most deaths during aging^4^. While DM is a metabolic disorder characterized by hyperglycemia and insufficiency in insulin production^5^, HT is a blood vessel disorder of persistent high tension^6^. In 2012 12.3% of the population in Mexico City suffered DM and 30.2% HT^7^; both conditions are independent, but can coexist and have a multiplying effect on complications, the frequency of occurrence is from 20 to 60% in DM patients, so HT is considered to contribute to the development and progression of complications in DM patients^8^.

One of the associated mechanisms to aging and that has been also related to DM and HT independently is oxidative stress (OS). OS is defined as any disturbance in the balance of anti-oxidants and pro-oxidants by an excessive formation and insufficient elimination of highly reactive molecules such as oxygen (reactive oxygen species, ROS)^5^, by catabolic and anabolic processes, mainly through the electron transport chain in the mitochondrial complexes^9^. High levels of ROS can eventually cause cell death^5^ by affecting proteins, lipids, and even DNA^9^.

Aging has been related to accumulation of genetic damage either by genotoxic attacks from exogenous sources such as UV light, ionizing radiation of chemical products, or endogenous lesions caused by ROS^10^ that become hard to cope by an organism with a diminished ability to counter these effects^11^. Aging reduces the activity of anti-oxidant enzymes such as the superoxide dismutase (SOD), glutathione peroxidase, and nitric oxide (NO); however, in the presence of diseases such as HT there is an increase in lipid peroxidation and an even lower NO production.

An increase in ROS production and lipid peroxidation has been reported in patients with DM, where chronic damage by OS is associated to occurrence and complications of the disease^12^. Moreover, in patients with HT it is known that OS alters the function of the endothelium and has a direct effect on the vascular contraction, due to the reduction of NO^13^. The increase in OS generates other alterations in the regulatory system that affect hypertension, including the positive regulation of the renin-angiotensin-aldosterone system, the activation of the sympathetic nervous system, the disturbances of the cellular signalling of the G-protein, inflammation, and alteration of T-cells function^13^.

ROS and lipid peroxidation measurement in elder patients with DM and or HT will allow us to establish the participation that OS can have in this specific segment of the population, which are much more prone to suffer chronic diseases and have a harder time reaching system homeostasis.

## Materials and methods

### Subjects

A transversal comparative study was performed with member from the Cohort of Obesity, Sarcopenia and Frailty of Older Mexican Adults (COSFOMA), a representative sample of older adults (>60 years of age), members of the Instituto Mexicano del Seguro Social (IMSS) of Mexico City^14^. One hundred and fifty four patients were recruited for the cohort follow-up of 2015 with DM and HT, DM, HT or without DM or HT. Only subjects that signed the consent letter according to the General Health Regulation, General Health Law, IMSS Health Regulation, and the Helsinki Declaration, participated in this study. The National Committee of Scientific Research as well as the Ethics Committee for Health Research IMSS approved this protocol (R-2015-785-022).

### Evaluation of reactive oxygen species (ROS)

ROS was measured by fluorometry, 5 μl of the serum sample and 85 μl of PBS 1X buffer was added to a 10 μl solution of diclorofluoresceína-diacetate (DCFDH-DA) to 15 μm (diluted in methanol), incubated in darkness for 30 minutes at 37 °C in an ELISA plate. The oxidation of the 2-7-diclorofluorescína (DCFH) to the fluorescent compound 2-7-diclorofluoresceína oxidized (DCF) by the presence of hydrogen peroxide, allows to use this reaction as a specific indicator of the formation of reactive species. The samples were then read by the fluorescence produced by the DCF 498 nm excitation at a 522 nm emission in a Cytation™ 5 Cell Imaging Multi-Mode Reader, previously calibrated with a standard curve.

### Determination of lipid peroxidation

The main reactive substance TBA is the malondialdehyde (MDA), a highly reactive three carbon dialdehyde, generated as one of the main bioproducts of the peroxidation of polyunsaturated fatty acids and is detected by spectrophotometry, the velocity of this reaction depends on the concentration of thiobarbituric acid (TBA), temperature and PH, the pigment generated has a maximum peak absorbance at 532-535 nm.

For this procedure 50 μl of serum were added to 25 μl of PBS in a 1x concentration, and 50 μl of the TBA reagent. Samples were placed in a boiling bath at 94°C for 20 minutes, then centrifuged at 10,500 rpm for 15 min, the supernatant was loaded into wells of an ELISA plate and read at 532 nm with an EPOCH spectrophotometer, previously calibrated with a stantard curve.

### Statistics

A descriptive analysis with measurements of central tendency and dispersion, as well as an inferential analysis of the values was obtained to determine the normality of the data with the D’Agostino and Pearsons test. According to which a non-parametric test was used for the comparison between two groups (Mann-Whitney U test) or more (Kruskal-Wallis test). A value of p<0.05 was considered significant. Correlation was measured using Spearman r.

## Results

Subjects were grouped according to the diseases studied: DM (Diabetes Mellitus), HT (Arterial Hypertension), DM and HT, and Control group (no DM or HT). When comparing reactive oxygen species (ROS) between groups, significant differences were found between the HT and the DM and HT groups (p<0.05); patients with HT presented a higher median compared to patients with both pathologies (Me=58.31 and Me=27.76 respectively) as can be seen in figure 1A.

**Figure 1.**
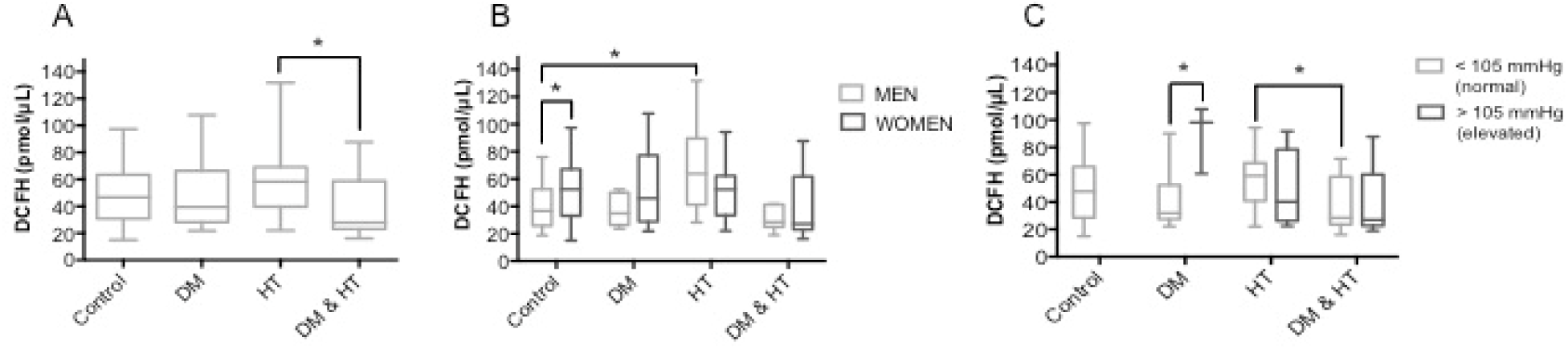
Reactive oxygen species (ROS), values are expressed as DCFH pmol/μl A. Between groups, control (without DM or HT), with DM, with HT and with DM and HT. B. Divided by sex, men in light grey and women in dark grey. C. Compared according to their mean arterial pressure; normal MAP in light grey and light MAO in dark grey. Significant differences *p<0.05.

When analysing data using gender as a variable significant differences in ROS measurements were obtain between men and women from the control group (p<0.05); men with a Me=36.69 and women Me=52.69. Also, men with HT and men in the control group had a significantly different ROS median (p<0.05; Me=63.85 and Me=36.69 respectively) (Figure 1B).

We also approached the data separating the groups according to the mean arterial pressure (MAP) between normal MAP (<105mmHg) and high MAP (>105mmHg) using the standard values used for clinical analysis. For ROS values significant differences (p<0.05) were found between HT patients and patients with both pathologies (DM and HT) and normal MAP. Also, the group of DM with elevated MAP, has significantly higher ROS values than the group with DM and normal MAP (figure 1C). Groups with elevated MAP showed no significant differences.

When analysing values of lipid peroxidation, (figure 2) significant differences were found only between sexes in the group with both pathologies (DM and HT) where men showed a Me=62.78 and women Me=28.86. No other changes were found for lipid peroxidation.

**Figure 2.**
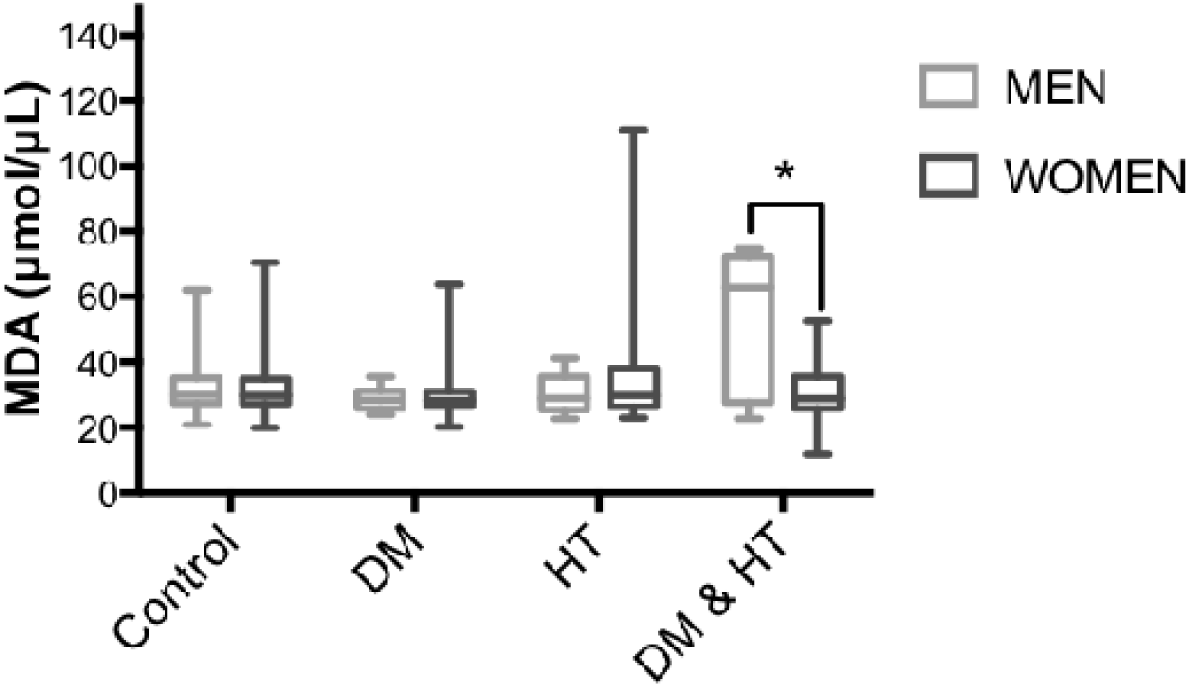
MDA expressed in μmol/μl divided by sex, light grey bars represent men and dark grey bars represent women. Significant differences were observed between men and women from the DM&HT group. Significant differences *p<0.05.

Finally, and since differences were observed amongst hypertensive patients, we decided to evaluate the association between blood pressure and both oxidative stress measurements. No correlation was found between blood pressure and ROS (r=0.1410) or lipid peroxidation (r=0.0939) as shown on figures 3A and 3B respectively.

**Figure 3.**
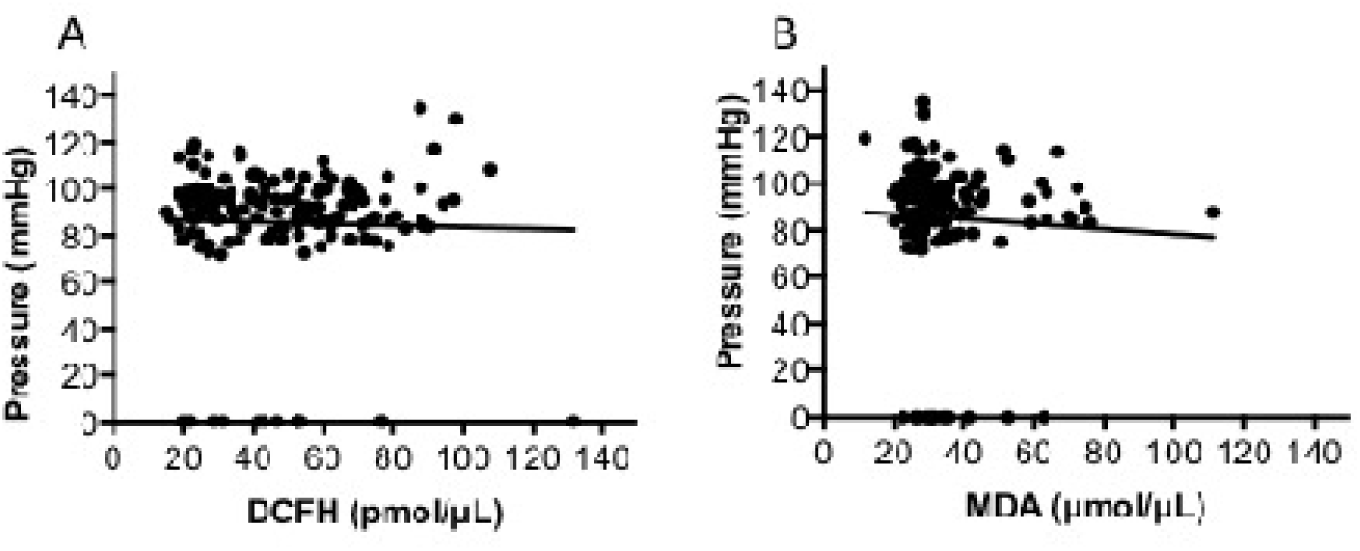
Association between blood pressure and oxidative stress. ROS measured by DCFH and lipid peroxidation by MDA. No correlation was found between pressure and oxidative stress.

## Discussion

Previous work has reported a raise in oxidative stress levels in patients with DM and H independently^15^. However, comorbidity of these diseases had not been studied, even less in an elderly population, a factor that has also been found to contribute to the increment in oxidative stress. Hence, this study focused on the changes in pro-oxidant agents of an elder population with DM, HT or both. When comparing both pathologies separately and combined in independent groups of older adults, we found that HT is the factor that contributes the most to the generation of reactive oxygen species (ROS).

On a first analysis of the data, patients with HA were found to generate more ROS than patients with DM and HA, which called our attention since we thought that the group with comorbidity would be the most affected. On a second analysis by gender, we observed that men with HT have more ROS than men in the control group. Additionally, significant differences were found between men and women with DM and HA, being men the most affected by oxidative stress.

Finally, even though all patients included in this study are users of the medical services of IMSS and should have some degree of control over their HA, we separated patients taking into account the mean arterial pressure. The group with normal pressure but HA still showed elevated ROS compared with controls. Hence, even though the HA is controlled, the disease has an effect at a molecular level (increased oxidative stress).

On the matter, the pathway were reactive oxygen species are generated in cardiovascular diseases such as HA, is through enzymatic systems such as the nicotinamide adenine dinucleotide phosphate (NADPH), the mitochondria complex, and under certain circumstances the nitric oxide synthase. More specifically, the NADPH can be activated by angiotensine II, a hormone that causes vasoconstriction and increases arterial pressure promoting oxidative stress in HA^16^.

The aforementioned results lead us to suggest that for the aged population studied here, HA is the most contributing factor to the generation of ROS. A controversial conclusion since in the beginning we thought that comorbidity of HA and DM would be the most significant component. Nonetheless, after reviewing the treatment used for DM patients, we found that most of them include anti-oxidants^17^. Also, it is known that certain drugs used for the treatment of HA interact with the antioxidant enzymes like superoxide dismutase (SOD) and catalases (CAT); patients with no HA treatment have a reduced activity of these enzymes, while patients receiving treatment for hypertension show an increased activity of both^13^.

On that matter, we must remember that the definition of oxidative stress is precisely an unbalance between anti-oxidant enzymes and pro-oxidant agents^5^; henceforth, the continuous consumption of these drugs could be affecting the increase in oxidative stress previously reported for DM patients. This, together with the anti-oxidants received by patients with DM and HA could contribute to a decrease in OS levels in patients with comorbidity.

We must emphasize that our population is exclusively of older adults, a group that has been described to have a reduction in anti-oxidants, which contributes to the increment in OE in the elders^11^. Since this is a cohort, were simultaneous studies are done, we were unable to evaluate these specific enzymes and can only discuss our results relative to the pro-oxidation of this complex system.

Within the analysis we performed by gender, we found relevant data. Women from the control group (no DM and no HA) of older adults generate the most ROS when compared with men of the same group. On the matter, contradictory results are found in the literature; some show an elevated OS in men than women^18^, and others determine an incremented OS in women^19^ or even no changes at all between gender^20^.

It is important to note that every study uses a different method to measure changes in oxidative stress. For example, in the study were men had greater OS, they measured the product of lipid peroxidation and changes in the anti-oxidative enzymes and found that along men’s life the first increments and the second diminishes compared to women^18^. On the other hand, when concluding that women have more OS, they measure reactive oxygen metabolites^19^.

In this study we found differences when measuring ROS, but did not find significance when analysing changes in lipid peroxidation. The women in this study are post-menopausal, a critical stage in life where endocrine changes occur caused mainly by the reduced production of estrogen with the capacity of working as anti-oxidants inhibiting or neutralizing the production of ROS; also, most of the symptoms after menopause include insomnia, anxiety and bad temper, all considered to be pro-oxidants^21^.

Independently of no accumulative effect being found when both pathologies occur at the same time, the general analysis of our data shows no changes at all in patients with DM. It has been suggested that physiological changes, including the increment of oxidative stress caused by DM and other chronic diseases, originate a phenomenon known as hormesis^22^. Hormesis is the process by which biological systems react to the exposition of chemicals, toxins, radiation and reactive oxygen species when these are modulated by aging and is reflected by a greater adaptation capacity and a better tolerance to stressing factors^23^. This could be the case for DM and the increase in blood glucose that in turn favours an increase in oxidative stress.

On the matter, a study were lipid peroxidation is measured by MDA in young and old adults with DM, found higher levels of oxidative stress in the young adults suggesting that it could be related to the time of diagnosis; meaning that elders gradually compensated the physiological changes derived from DM through hormesis^22^ reflecting it as lower OS.

On the other hand, clinical diagnosis of DM is more frequent in people between 40 and 50 years of age. Patients with little care of the illness that develop severe DM or DM with complications are more likely to die before reaching 60, the youngest patients used for this study. Hence, we must consider that our population with DM may be biased. However, this is the case for the general population, but we consider our results to be a reflection of what happens in an aged population with DM.

Finally, we found no correlation between blood pressure and ROS or lipid peroxidation. Again, it seems as if HA defines OS in this population, independent of the current state of the patient; in this case, the measurement of MAP on the day the sample was taken. HA doesn’t seem to have compensation by hormesis but an aggregative damage by OS.

In our study, HA is the most contributing factor in the production of reactive oxygen species. The aged population is without a doubt a hard group to evaluate due to the effect of aging itself over biological factors. Hence, this subject can be evaluated incorporating more information regarding other factors that are known to influence oxidative stress such as quality of life in our population.

## Conclusion

In a population from Mexico City, healthy elder women have a higher ROS production probably due to post-menopause. On the other hand, HA is the most contributing factor to an increase in ROS production while DM does not affect either measurement of oxidative stress (ROS or lipid peroxidation).

## Acknowledgments

We thank the technical support of Ana Laura Colin Gonzalez and Abel Santamaría del Ángel. This project was done with the participation of the Collaborative Group for the Study and Management of Dementia of the Instituto Mexicano del Seguro Social.

